# Neural Correlates of Different Randomization Tasks

**DOI:** 10.1101/2024.03.27.586961

**Authors:** Maja Guseva, Carsten Bogler, Carsten Allefeld, Ece Büşra Ziya, John-Dylan Haynes

## Abstract

In some cases, when we are making decisions, the available choices can appear to be equivalent. When this happens, our choices appear not to be constrained by external factors and instead we can believe to be selecting “randomly”. Furthermore, randomness is sometimes even explicitly required by task conditions such as in random sequence generation (RSG) tasks. This is a challenging task that involves the coordination of multiple cognitive processes, which can include the inhibition of habitual choice patterns and monitoring of the running choice sequence.

It has been shown that random choices are strongly influenced by the way they are instructed. This raises the question whether the brain mechanisms underlying random selection also differ between different task instructions. To assess this, we measured brain activity while participants were engaging in three different variations of a sequence generation task: Based on previous work, participants were instructed to either (1) “generate a random sequence of choices”, (2) “simulate a fair coin toss”, or (3) “choose freely”.

Our results reveal a consistent frontoparietal activation pattern that is shared across all tasks. Specifically, increased activity was observed in bilateral inferior and right middle frontal gyrus, left pre-supplementary motor area, bilateral inferior parietal lobules and portions of anterior insular cortex in both hemispheres. Activity in the mental coin toss condition was higher in right dorsolateral prefrontal cortex, left (pre-) supplementary motor area, a portion of right inferior frontal gyrus, bilateral superior parietal lobules and bilateral anterior insula. Additionally, our multivariate analysis revealed a distinct region in the right frontal pole to be predictive of the outcome of choices, but only when randomness was explicitly instructed.

These results emphasize that different randomization tasks involve both shared and unique neural mechanisms. Thus, even seemingly similar randomization behavior can be produced by different neural pathways.

## Introduction

The ability to elicit seemingly random behavior can be a very useful in many different situations and has been observed across such diverse domains as predator evasion (Humphries & Driver, 1970), exploration (Wilson et al., 2014), creativity (Benedek et al., 2012), improvisation (de Manzano & Ullén, 2012) and decision-making (Icard, 2019). Absence of randomization ability, which can go hand in hand with behavioral stereotypy, has been observed in various psychopathologies (Horne et al., 1982).

Human randomization behavior has been commonly studied using random sequence generation (RSG) tasks (Daniels et al., 2003; Guseva et al., 2023; Heuer et al., 2010; Jahanshahi et al., 2000; Naefgen & Janczyk, 2018; Nickerson, 2002; Peters et al., 2007; Schneider et al., 2004). In such tasks participants make consecutive choices between different choice option sets in a seemingly random way (see Nickerson, 2002). RSG is a complex task that requires the coordination of different higher-order cognitive operations. These include updating and monitoring working memory to maintain and retrieve previous choices as well as inhibition of overlearned, “patterned” responses (e.g., ascending or descending the number line, Jahanshahi et al., 2006). Dysfunctions in these cognitive functions can be due to transient executive disruptions (Heuer et al., 2005; Jahanshahi et al., 1998; Naefgen & Janczyk, 2018) or psychopathological deficits (see Horne et al., 1982 for a review) and can result in common choice biases, such as repetition avoidance, cycling bias or seriation bias (Peters et al., 2007).

### Neural bases of RSG

Compared to the breadth of studies examining various facets of executive function, such as Wisconsin Card Sorting Test (Yuan & Raz, 2014), Tower of Hanoi (Anderson et al., 2005) or Stroop task (Laird et al., 2005), the neural mechanisms of randomization behavior have only rarely been studied. Work by Jahanshahi and colleagues using transcranial magnetic stimulation (TMS) and positron emission tomography (PET) highlighted the role of the left DLPFC in an RSG task using letters as a choice set (Jahanshahi et al., 1998, 2000; Jahanshahi & Dirnberger, 1999). The authors suggested that the left DLPFC stops the spread of activation in the number associative network in the superior temporal cortex, thereby preventing prepotent responses, such as habitual counting. Other identified areas of activity include anterior cingulate cortex, premotor cortex and superior parietal lobules. Similarly, using near-infrared spectroscopy (fNIRS) Koike et al. (2011) showed significantly increased oxygenated hemoglobin in bilateral DLPFC and ventrolateral prefrontal cortex (VLPFC). A few other studies corroborated the involvement of DLPFC, with repetitive TMS (Knoch et al., 2004) and electroencephalography (EEG), presenting evidence of left frontal negativity during RSG (Joppich et al., 2004; Schneider et al., 2004).

Daniels et al. (2003) challenged the role of the DLPFC as a central node in RSG. At slower response rates (1Hz), they observed an increased activation of bilateral DLPFC, premotor cortex, anterior cingulate, inferior and superior parietal cortex and cerebellar hemispheres, while faster rates (2 Hz) led to reduced activation in these areas. Whereas Jahanshahi et al. (2000) reported a production rate dependent decrease specifically in bilateral DLPFC and superior/inferior parietal cortex, Daniels et al. (2003) observed a more general decrease and proposed that RSG is subserved by multiple regions in a network instead of only the DLPFC.

In summary, the findings converge towards the idea that the dorsolateral activation reflects some sort of suppression mechanism of habitual responses. While the DLPFC seems to play an important role in the task, it is likely a part of a distributed network of brain areas that work together to generate randomness, potentially working in unison with parietal, cingulate and cerebellar cortices (Artiges et al., 2000; Daniels et al., 2003).

### Task diversity in RSG studies

There is no standardized experimental procedure in RSG studies. Instead RSG has employed a variety of tasks and parameters, such as different numbers of choice options, such as the digits from 0-9 (Joppich et al., 2004), binary sets e.g., 0/1 or heads/tails (Nickerson & Butler, 2009), the letters A-I (Jahanshahi & Dirnberger, 1999), or nouns (Heuer et al., 2010). Another variable task parameter across RSG experiments is the method of instructing participants to be random, which ranges from analogies like drawing numbers from a hat (Jahanshahi et al., 2000), tossing a coin (Nickerson & Butler, 2009) or roll a die (Knoch et al., 2004), via instructing to avoid specific patterns (Azouvi et al., 1996; Daniels et al., 2003) all the way to emphasizing unpredictability (Finke, 1984).

In a recent study we confirmed that randomization performance depends on the precise instructions (Guseva et al., 2023). In that experiment, the requirement to choose randomly was explicitly stated in some instruction conditions (“choose randomly”) while being more indirectly implied in others. For example, participants were instructed to choose randomly, choose freely, or even select the darker of two coins, despite the fact that unknown to the participants they were equally bright. We found that the instructions to “mimic a fair coin toss” and to elicit an “irregular pattern” led to the most random sequences (as quantified by conditional entropy, see below). Building upon this behavioral paradigm, we conducted an fMRI study with 85 participants to investigate the underlying neural substrates behind the different methods of inducing randomness.

### The present study

In the present exploratory study, we investigated the neural bases of random sequence generation under different randomization instructions. Participants were required to make repeated binary choices between heads and tails of a coin while we acquired functional magnetic resonance imaging data from their brain activity. Importantly, we used three task variations that elicit differences in randomization behavior based on a previous study (Guseva et al., 2023). The Explicit Randomness (ER) task group received the instruction to “*make random choices*”, the Free Choice group (FC) task group to “*choose freely between the two sides*”, and the Mental Coin Toss (MC) task group to “*simulate the output of a fair coin toss*”. These tasks were chosen based on previous behavioral work (Guseva et al., 2023) and because they present conceptually different ways to elicit random behavior. In line with that previous work, we expected significant differences in randomization measures between MC sequences compared to ER and FC and no differences between ER and FC.

Previous research suggests shared cognitive processes and neural substrates between free choice tasks and RSG (Naefgen & Janczyk, 2018). Hence, we expected a frontoparietal pattern of activity across all three tasks. Specifically, we expected activity in the DLPFC, inferior and superior frontal gyri, premotor as well as inferior and superior parietal cortices. Additionally, based on the reviewed research results we hypothesized the recruitment of anterior cingulate cortex and anterior insula in all tasks. These expectations were not part of the preregistration.

Tasks like ER and MC likely require the coordination of multiple cognitive processes, such as choice history monitoring, inhibition of patterned responses and holding task-related information in working memory. For this reason, we anticipated an increased activation of control-related regions in ER and MC. In contrast, we presumed that most of these higher-order processes were less relevant for completing an FC task. In this task we expected the fluctuation between task-related and task-unrelated (mind-wandering) activity to be highest, which might be subserved by default mode network (DMN), i.e. posterior cingulate cortex and medial prefrontal cortex (Mittner et al., 2014), and mediating salience network activity, involving anterior cingulate cortex and anterior insula activation (Schimmelpfennig et al., 2023). Lastly, during the mental simulation of a coin toss, we predicted increased posterior parietal activation, particularly in the precuneus (Cavanna & Trimble, 2006), reflecting potential involvement in visual imagery.

## Methods

As in our previous behavioral study (Guseva et al., 2023), we studied the neural effects of different randomization task instructions in a between-subjects design to avoid carry-overs between tasks. Each participant received one of the three task instructions: Explicit Randomness (ER), Free Choice (FC) and Mental Coin Toss (MC). In ER we explicitly stated to choose the sides of the coins in a random way: “*You have to choose the sides of the coin randomly*”. In FC the instruction was to choose freely between the coins without any constraints: “*There is no right or wrong answer, we want you to decide spontaneously. Which side you choose in every trial is your own free choice*”. In MC the task was to choose according to a coin toss: “*You have to simulate a coin toss in your head and choose the side that came up. The goal is to produce a sequence of choices that is not different from the results of a real fair coin toss.*”

### Procedure

Prior to the experiment, each invited participant gave their informed consent of participation by signing a consent form. Next, participants completed a training session outside the scanner to familiarize themselves with the task. After that, participants entered the MRI scanner and began the session with an anatomical scan, during which the instructions and training sessions were repeated to practice the choice selection using the response button boxes. During the main experiment, participants completed 6 functional runs in total, each run consisting of 3 experimental blocks (Fig. 1A). One experimental block comprised 30 trials and had a duration of 90 s. The first two blocks were separated by a 20 s break. The break following the third block was 30 s, after which two Likert-scaled questions were administered (see below).

**Figure 1.**
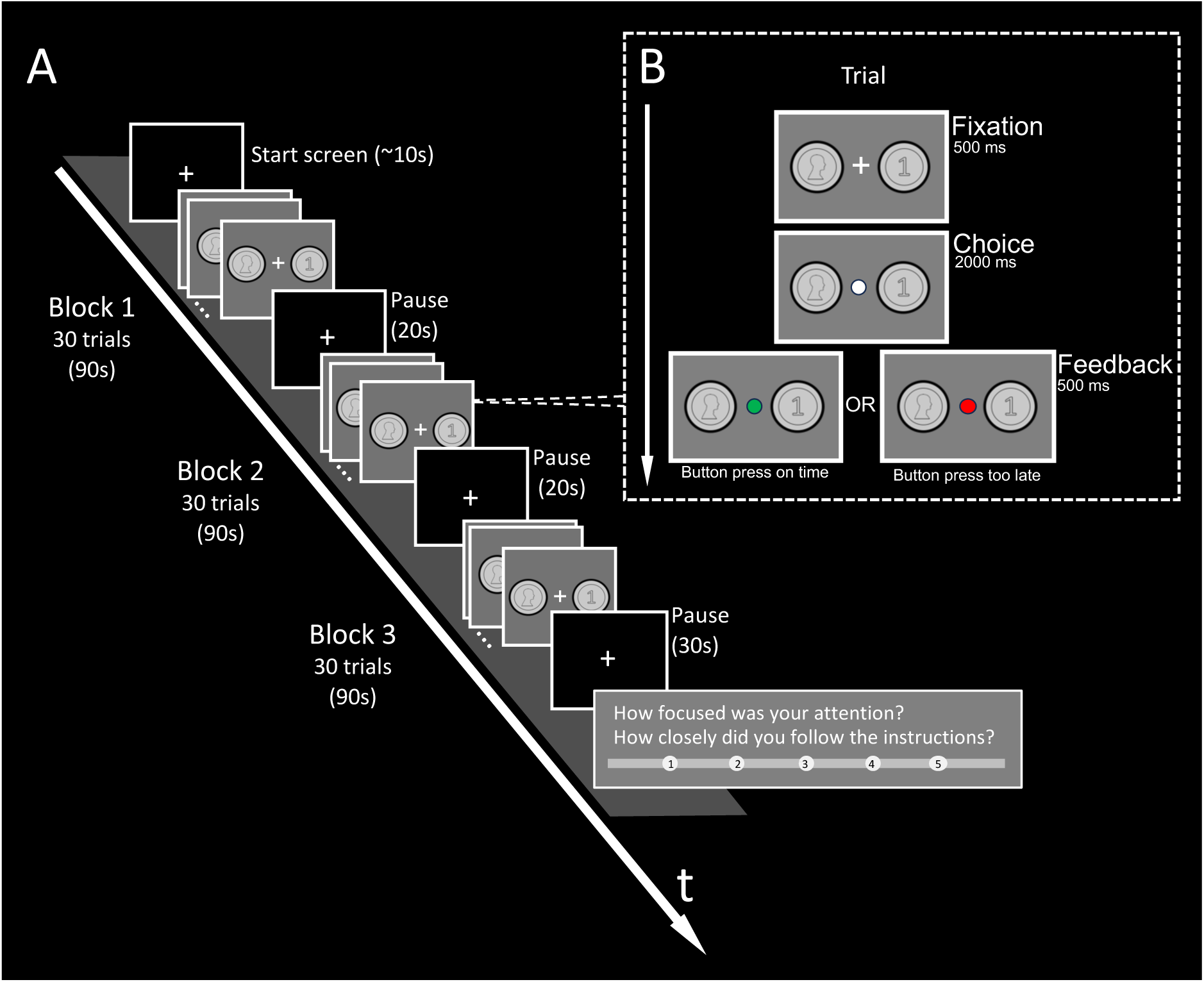
Experimental Procedure. A: Procedure of a single experimental run. The study comprised 6 functional runs, each consisting of 3 experimental blocks of 30 trials each. Short breaks of 20s and 30s separated the blocks. At the end of each run, participants provided responses on a five-point Likert scale regarding their focus and instruction adherence. B: Procedure of a single experimental trial. Following a fixation period of 500ms, the participant was required to make a choice within 2000ms for one of the two stimuli and indicate their choice with a button press using either the left or the right hand with a button box. During the subsequent feedback phase (500ms), the circle turned green if the response was given within the allotted time window and red if it was given too late.

A trial consisted of three stages (see Fig. 1B). First, participants fixated on a cross in the center of the screen (500ms). On the left and right of the fixation cross images of the heads and tails sides of a coin were presented. The position of either side of the coin was determined randomly in the beginning of each participant’s session and didn’t change throughout the experiment within subject. Next, the fixation cross turned into a white circle indicating that a choice between heads or tails using a button press was required (2000 ms). If the button was pressed within the allotted 2000 ms, then the circle filled with green color (successful trial) and if not, with red color (failed trial) for 500 ms.

At the end of each run, we asked participants to answer two questions on a five-point Likert scale: “How focused was your attention during this run?” (1 = ”Not focused at all” – 5 = ”Very focused) and “How closely did you follow the instructions during this run?” (1 = ”Did not follow at all”, 5 = ”Followed very closely”). Before starting the next run the participants could take a break for as long as they needed. At the end of the experiment, participants were required to write a free text about their strategy during the task (outside of the scanner).

Participation in the experiment was compensated with 12 € per hour. The whole experiment took on average 1.5 h per participant. The behavioral task was implemented in PsychoPy (v2022.2.4, https://www.psychopy.org/) (Peirce et al., 2019). The experimental procedure was approved by the Ethics Committee of the Department of Psychology (Humboldt University of Berlin). The experimental design and analyses were preregistered at https://osf.io/m7y92.

### Participants

A total sample of 90 neurologically healthy, right-handed participants (54 females) with normal or corrected-to-normal vision and who fulfilled the standard MRI safety criteria was acquired. Previous neurocognitive studies of RSG had sample sizes ranging from 6 to 18 (Daniels et al., 2003; Jahanshahi et al., 1998, 2000; Joppich et al., 2004; Knoch et al., 2004; Schneider et al., 2004). Given our between-subjects design, we decided to double the sample size to 30 in each of the three groups. Participants were recruited via participant mailing lists and flyers on campus. Each participant was allocated to one of the three conditions such that mean age and sex were balanced between the groups. This was achieved by an adaptive stratified sampling method. The script of this sampling method can be found this repository: https://github.com/m-guseva/balanced-group-assignment.

From the initial sample of 90 participants, we had to exclude 5 participants because they reported not following the task instructions (3 from the ER group, 1 from the FC group, and 1 from the MC group). The final sample contains 85 datasets. The age distribution ranged from 19 to 36 years and was as follows: ER: M=25.9, SD=4.4 (n=27); FC: M=26.8, SD=4.2 (n=29); MC: M=25.5, SD=3.7 (n=29). The sex distribution per condition is 17 females in ER, 18 females in FC and 18 females in MC.

### Image acquisition

The images were acquired on a 3T SIEMENS MAGNETOM Prisma fit MRI scanner (Siemens, Erlangen) with a 64-channel phased array head coil at the Center for Cognitive Neuroscience Berlin. At the beginning of the scanning session a T1-weighted 3D MPRAGE whole-brain anatomical image was acquired (TR = 1930 ms, TE = 3.52 ms, FOV = 20.5 cm, flip angle = 8°, voxel size: 0.8 mm, 208 slices (ascending), acquisition time 5:39 min). The functional imaging data was collected using gradient-echo echo planar T2*-weighted imaging (EPI-Factor = 80, TR = 1500ms, TE = 33ms, FOV: 20cm, flip angle = 70°, voxel size: 2.5mm, slices = 54 (interleaved), acquisition time 5:53min) with A/P phase-encoding direction. In total, we obtained six runs of whole-brain functional data from each participant, with 230 volumes and a duration of 5min 53s per run. Additionally, field maps were acquired for later use in distortion correction in the preprocessing pipeline (TR = 400ms, TE1 = 4.92ms, TE2 = 7.38ms, flip angle = 60°, acquisition time = 55s). The stimuli were displayed on a screen using an LCD projector (1920 x 1080, 120Hz frame rate). The participants were able to view the projection through a mirror attached to the head coil. The choices were logged via two 2-button response boxes in each hand.

### Data analysis

#### Behavioral Analysis

Based on the recorded sequence of button presses we assessed the degree of randomness using a procedure established previously (Guseva et al., 2023). This method consists of modeling the sequences of discrete choices as a Markov chain. Here we determined the optimal Markov order (the order k that describes the number of past choices t-k that determine the present choice at t) which reflects the temporal extent of sequential dependencies in the sequence. The transition probabilities Pr(X_t_| X_t-1_, X_t-2_, …, X_t-k_) at k=3 were used to calculate conditional entropy (see Guseva et al. (2023) for a detailed description of the method). Both randomness measures were calculated for a whole run and then averaged across the runs to get one value per person, using the mean for conditional entropy and median for the optimal Markov order. Following the procedure in Guseva et al. (2023), we used the non-parametric Kruskal-Wallis test at a significance level of 0.05 to detect differences in medians between task conditions.

Also similar to our previous study, we determined the run lengths and proportion values of the sequences. The run lengths, proportion values and reaction times (i.e., the duration between the start of the choice phase and the button press) were determined within a block, then averaged across blocks and then averaged across all runs to get one summary value per person. We compared these summary values between conditions using ANOVA, followed by post hoc tests at a significance level set at 0.05 (Tukey-Kramer adjusted).

#### Imaging Analysis

The images were processed and analyzed using the Statistical Parametric Mapping toolbox SPM12 (Wellcome Trust Centre for Neuroimaging, Institute for Neurology, University College London, London, UK, https://www.fil.ion.ucl.ac.uk/spm/). Anatomical regions were labeled using the probabilistic cytoarchitectonic maps in the SPM Anatomy toolbox for MATLAB (Eickhoff et al., 2005) and FSL anatomical labels (*mni2atlas* toolbox, Mascali (2024). Visualizations of brain activations were created with MRIcroGL (Neuroimaging Tools & Resources Collaboratory https://www.nitrc.org/projects/mricrogl).

The images were unwarped using the fieldmaps entered into the fieldmaps toolbox of SPM and spatially realigned to the first image using an affine rigid-body transformation to correct for head motion. The functional and anatomical images were coregistered. We smoothed the images with a 6 x 6 x 6 mm^3^ FWHM Gaussian Kernel for the univariate analysis only. We normalized the images before entering into the second level analysis for both univariate and multivariate analyses to the standard Montreal Neurological Institute (MNI) space using unified segmentation (Ashburner & Friston, 2005).

We used a mass-univariate general linear model (GLM) to model the time series of the blood oxygenation level dependent (BOLD) signal in each voxel with the experimental block onset as the regressor (duration: 90s). A canonical hemodynamic response function (HRF) as implemented in SPM was used to convolve the regressors. A default high pass filter of 128s was used. The six motion parameters were included as additional nuisance regressors of no interest. At the second level we used a random effects approach to test for group differences between the three task variations ER, FC and MC. The ANOVA additionally included age, sex and responses to the instruction adherence and attention questions as predictors. All effects are reported at p<0.001; family-wise error (FWE) cluster-corrected at p<0.05.

For the multivariate analysis, we estimated a first level GLM with left/right button press onsets and motion parameters as regressors, this time based on unsmoothed data. The resulting parameter estimates of the choice regressors served as inputs for the MVPA implemented in The Decoding Toolbox (TDT) (Hebart et al., 2015). A linear support vector machine (SVM) was trained on parameter estimates of 5 runs and used to predict the button press on the left-out test data, resulting in prediction accuracy maps for each individual. Here we used leave-one-run-out cross-validated searchlight decoding with a radius of 4 voxels. The accuracy maps were then normalized and smoothed and entered into a second level ANOVA. Please note, that this analysis does not fall under the criticism of Allefeld et al. (2016), because accuracies were not compared between groups. Age, sex and responses to the instruction adherence and attention questions were used as covariates to test for differences between the three groups. This is an additional exploratory analysis and was not part of the preregistration.

## Results

### Behavioral Results

#### Focus and instruction adherence per condition

Fig. 2 shows the distribution of answers for each run. Overall, 63.53% of the participants answered that they were “very focused” to “focused” and 89.41% of people reported to have followed the instructions “very closely” to “closely”. The ratings in terms of focus decreased up until the third run and then slightly increased again whereas instruction adherence seems to be more stable over the runs. We did not find a significant difference between conditions for both focus and instruction adherence ratings (𝜒^!^=4.89, p=0.09 and 𝜒^!^=1.32, p=0.52 respectively).

**Figure 2.**
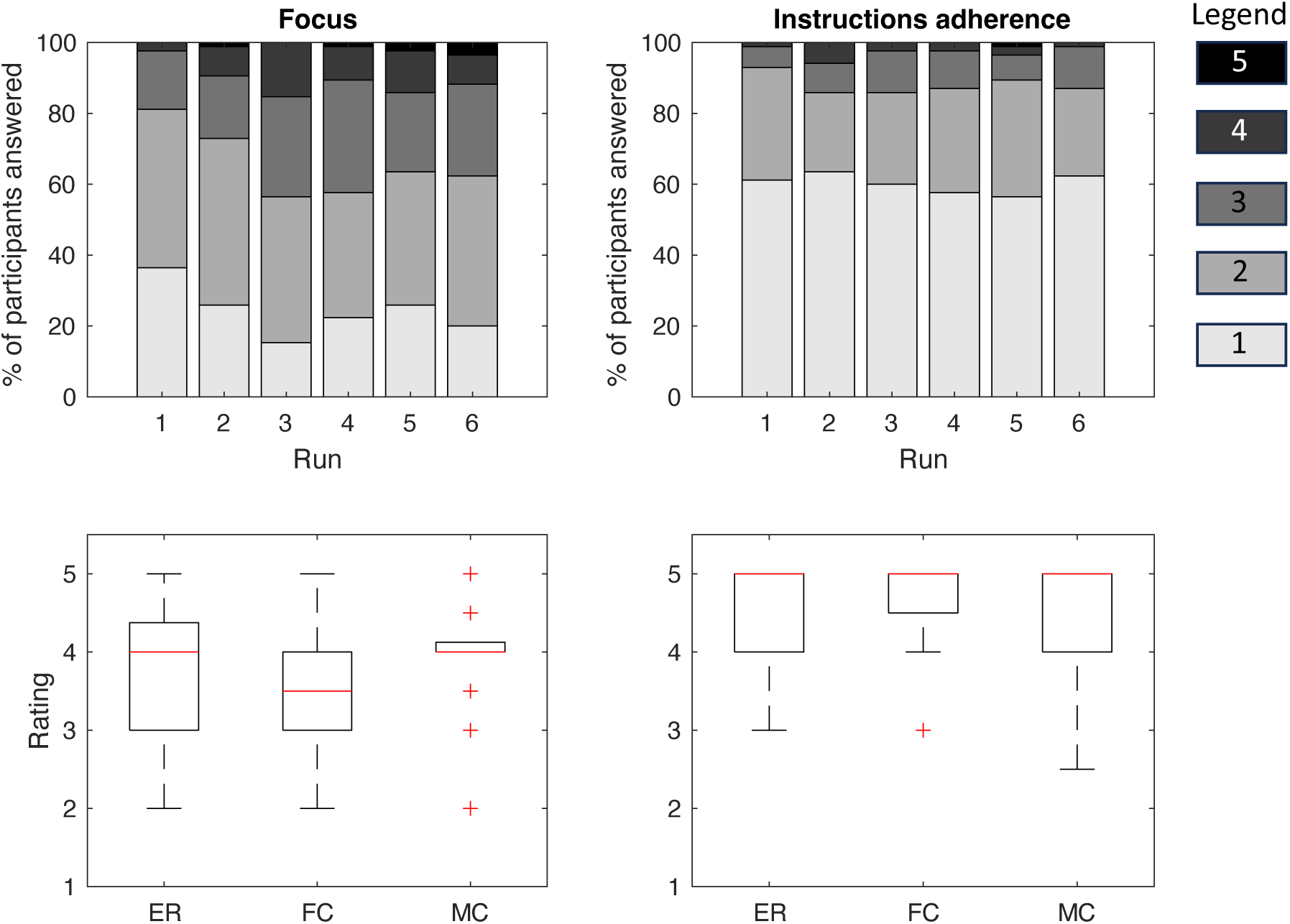
Distribution of responses per condition. Results of focus and instruction adherence questions on a 5-point Likert scale from 1 = ’Very focused’/Followed very closely’ to 5 = ’Not focused at all’/’Did not follow at all’. Top: Percentage of participants per rating in each run, Bottom: Boxplots showing the distribution of median ratings across runs for each condition. *ER*: Explicit Randomness, *FC*: Free Choice, *MC*: Mental Coin Toss.

#### Proportion value, run length, reaction time

The average proportion value reflects the balance between choices of heads and tails, where 0.5 indicates equal frequency of both options and 1 indicates the choice of exclusively one option. This value was lowest in MC (*M*=0.55, SD=0.04), followed by ER (*M*=0.56, *SD*=0.03). FC had the least balanced sequences on average (*M*=0.58, *SD*=0.07). A one-way ANOVA revealed a significant difference between the groups (F(2,82) = 3.34, p=0.04). The follow-up t-tests adjusted for multiple comparisons with Tukey-Kramer test showed that FC was significantly different from MC (p=0.03).

The average run lengths per condition are shown in Fig. 3. The one-way ANOVA revealed no significant difference between the groups (F(2,82) = 2.31, p=0.1). The mean length of runs clustered around 2 (ER: *M*=2.01, *SD*=1.15; FC: *M*=2.25, *SD*=1.15; MC: *M*=1.71, *SD*=0.39).

**Figure 3.**
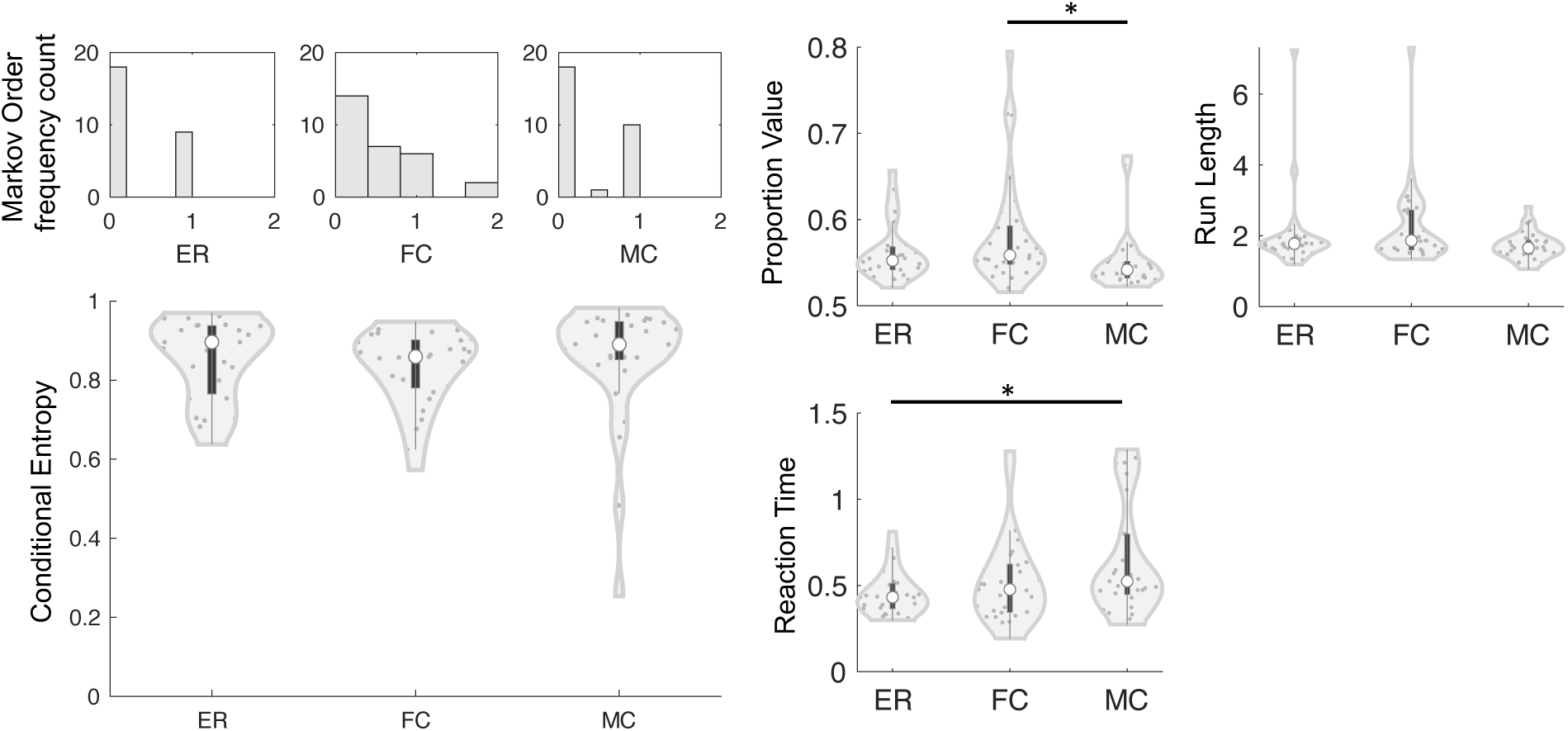
Behavioral results. Distribution of sequence properties in each condition. The histograms show the frequency count of orders per condition. In the violin plots, each dot represents the averaged value for one subject. The black bars in each violin plot are boxplots and indicate the interquartile range with the median marked by a white dot, the outline of each violin represents the kernel density estimate. Left: Randomness measures conditional entropy and Markov order were calculated for an entire run and then averaged across all runs using the mean for the conditional entropy and median for the Markov order. Right: Proportion value, run length and reaction times were determined for a block and then averaged over blocks and runs using the mean. n=85 in each metric plot, * p<0.05.

In terms of reaction times, a one-way ANOVA revealed a significant difference between conditions (F(2,82)=4.22, p=0.02). A post-hoc test showed that the reaction times between ER (*M*=0.45, *SD*=0.13) and MC (*M*=0.64, *SD*=0.31) differed significantly (p=0.014). FC (*M*=0.52, *SD*=0.25) did not differ significantly from the other groups (see Fig. 3).

#### Conditional Entropy and optimal Markov Order

Fig. 3 shows the distribution of individuals’ randomness measures, optimal Markov order and conditional entropy, averaged over runs. The conditional entropy values clustered around 0.90 in ER and MC (*Md*=0.9, *IQR*=0.17; and *Md*=0.90, *IQR*=0.10 respectively), whereas the value in FC was lower (*Md*=0.86, *IQR*=0.12), suggesting that ER and MC sequences displayed higher levels of randomness. The Markov order values were also similar in ER and MC with *Md*=0 and *IQR*=1 in both, but higher in FC (*Md*=0.5, *IQR*=1). As a benchmark a set of simulated pseudorandom sequences of same length using MATLAB’s default “Mersenne twister” algorithm of length 540 exhibits a median conditional entropy of 0.97 (*IQR*=0.01) and median Markov order of 0 (*IQR*=0). Neither Markov order values nor conditional entropy values differed between conditions (Kruskal-Wallis test, 𝜒^!^=0.9, p=0.64; 𝜒^!^=3.71, p=0.16 respectively).

### Imaging Results

#### Univariate conjunction analysis

We used a conjunction analysis (Friston et al., 2005) to identify activations that are shared across all three tasks (Fig. 4.B left; Fig. 6; table 1). Common activity was increased in a large bilateral frontal cluster consisting of regions in the inferior frontal gyri (IFG) and extending into portions of the anterior insula (aINS). Overlapping activations were also present in left premotor cortex/pre-supplementary motor area (pre-SMA) accompanied by smaller clusters involving bilateral precentral gyri and segments of middle frontal gyri. Additionally, BA44 (partially overlapping with Broca’s area) was activated. Our analysis revealed also increased activations in bilateral inferior parietal lobules (IPL), putamen, occipital cortex and cerebellum across all tasks.

**Figure 4.**
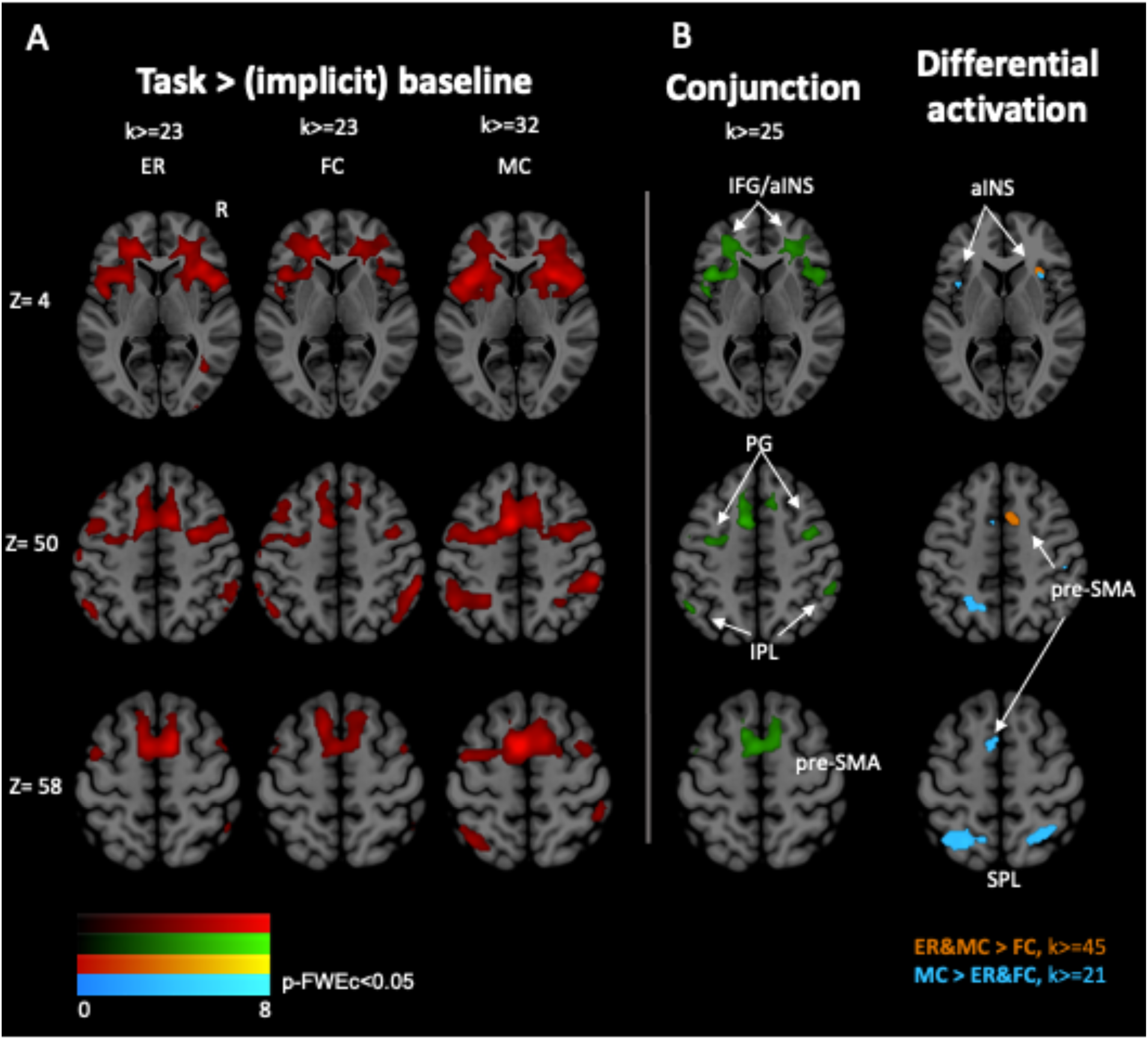
Whole-brain group level analysis (GLM). Results of the whole-brain group level analysis at three different transversal slices (Z=4, Z=50, Z=58). Shown are significantly activated clusters overlaid on the Montreal Neurological Institute space (MNI) template, p<0.001, FWE cluster corrected at p<0.05 (corresponding extent threshold k is displayed for each contrast). A: Contrasts of task activation against (implicit) baseline for each condition. B, left: conjunction analysis across all tasks. B, right: Differential activations for the contrasts MC>ER&FC and ER&MC>FC. Abbreviations: *ER*: Explicit Randomness, *FC*: Free Choice, *MC*: Mental Coin Toss, *IFG*: inferior frontal gyrus, *aINS*: anterior insula, *pre-SMA*: pre-supplementary motor area, *PG*: precentral gyrus, *IPL*: inferior parietal lobule, *SPL*: superior parietal lobule.

**Table 1.** MNI coordinates of peaks of whole-brain group level conjunction analysis results with p<0.001 uncorrected; family-wise error (FWE) cluster-corrected p<0.05 (k>=25). Abbreviations: Hem= Hemisphere, L=left, R=right; k=cluster size.

|  | Anatomical Area | Hem | t-value | k | x | y | z |
| --- | --- | --- | --- | --- | --- | --- | --- |
| Frontal | Inferior frontal gyrus | L | 5.57 | 1677 | -24 | 38 | 10 |
|  | Inferior frontal gyrus | R | 5.74 | 1575 | 30 | 32 | 10 |
|  | Pre-supplementary motor area | L | 5.67 | 1056 | -10 | 8 | 48 |
|  | Precentral/middle frontal gyrus | R | 4.62 | 116 | 38 | -4 | 50 |
|  | Premotor area BA6 | L | 4.29 | 189 | -34 | -8 | 48 |
|  | Broca's area BA44 | L | 4.25 | 71 | -56 | 8 | 22 |
| Parietal | Inferior parietal lobule | L | 4.66 | 66 | -50 | -56 | 50 |
|  |  | R | 4.34 | 114 | 58 | -40 | 46 |
| Cerebellum |  | R | 4.02 | 47 | 40 | -56 | -32 |
| Putamen |  | L | 4.65 | 102 | -26 | 4 | -4 |
|  |  | R | 4.02 | 28 | 26 | 4 | -4 |
| Occipital | Visual cortex V3 | L | 4.01 | 25 | -14 | -86 | -10 |

#### Univariate differential analysis

Next, we assessed any evidence for *differential* activity between conditions. Compared to ER and MC, activity in the right pre-SMA, right insula and left cerebellum activity was reduced in FC (Fig. 4.B right; table 2).

**Table 2.**
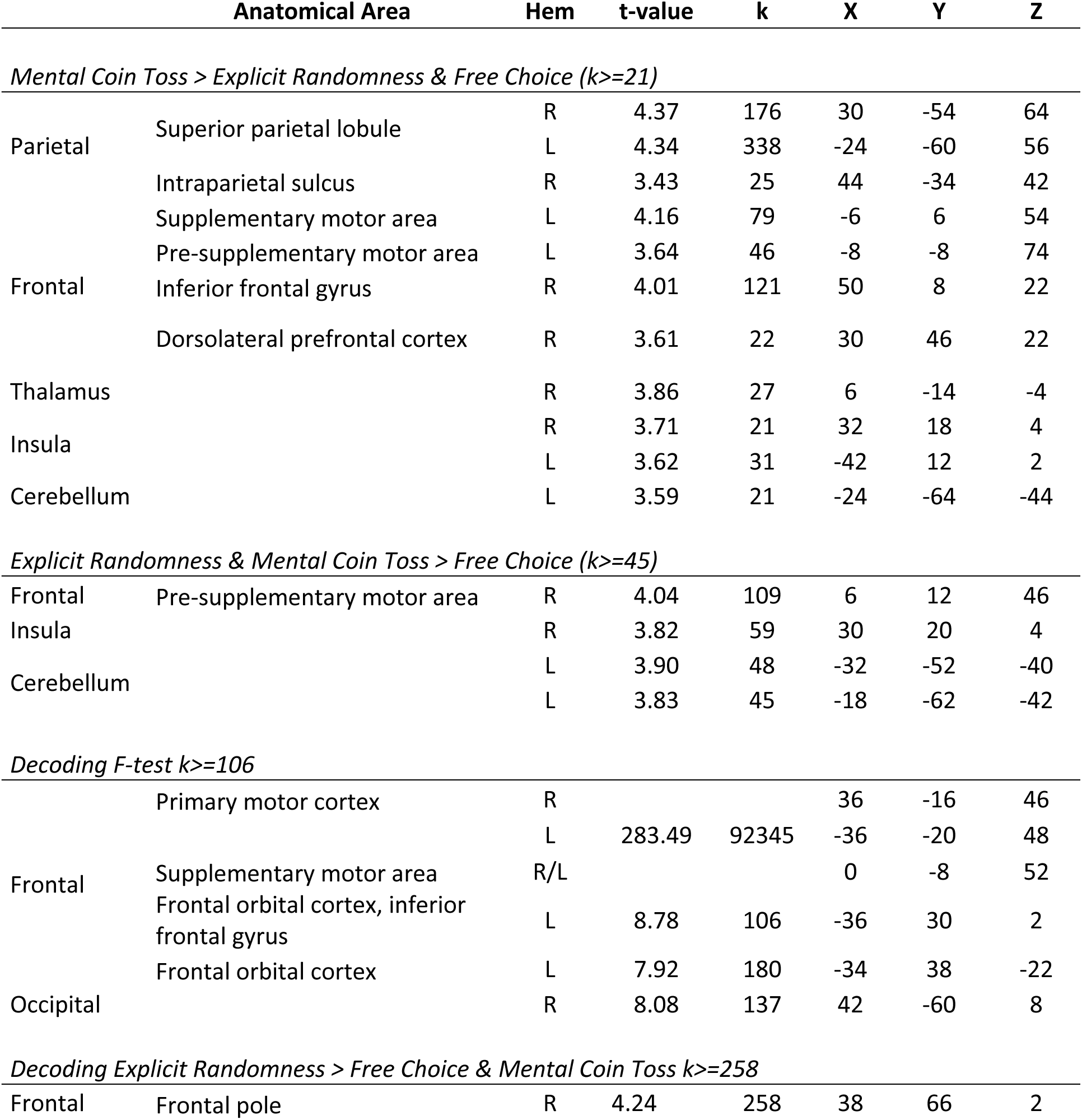
MNI peak coordinates of significantly activated clusters for different contrasts (whole-brain) p<0.001 uncorrected; family-wise error (FWE) cluster-corrected p<0.05 (k > 45) on the group level. Abbreviations: Hem= Hemisphere, L=left, R=right; k=cluster size. [add brain region abbreviations].

During the MC task, we observed a stronger bilateral frontoparietal pattern of activations compared to ER and FC combined (Fig 4.B right; table 2). In the frontal lobes, a subregion of the right DLPFC (Fig. 6) was more activated. More posteriorly, there was increased activity in right BA44, bordering on inferior frontal and precentral gyrus as well as two clusters in the left hemisphere comprising pre-SMA and SMA. The MC task also elicited stronger activation in bilateral portions of insular cortex along with regions in bilateral thalamus and left cerebellum compared to FC and ER combined. Our analysis further identified clusters of increased activity in the bilateral superior parietal lobules (SPL), and right intraparietal sulcus (IPS).

#### Multivariate analysis

Next, we used multivariate pattern analysis to identify brain regions from which we can predict subjects’ specific trial-by-trial choices. Fig. 5.A and table 2 show regions from where choices can be decoded across any of the three conditions. As expected, bilateral primary motor cortex areas show the highest decoding accuracy across all conditions. Additionally, areas in the left frontal lobes encompassing parts of IFG, frontal orbital cortex and insular cortex showed significant above-chance decoding accuracy. The comparison of accuracy values of ER versus FC and MC revealed significantly higher accuracy values in the right frontal pole (Fig 5.B; Fig 6; table 2). Figure 6 shows this cluster plotted together with the differential and shared univariate activations for comparison.

**Figure 5.**
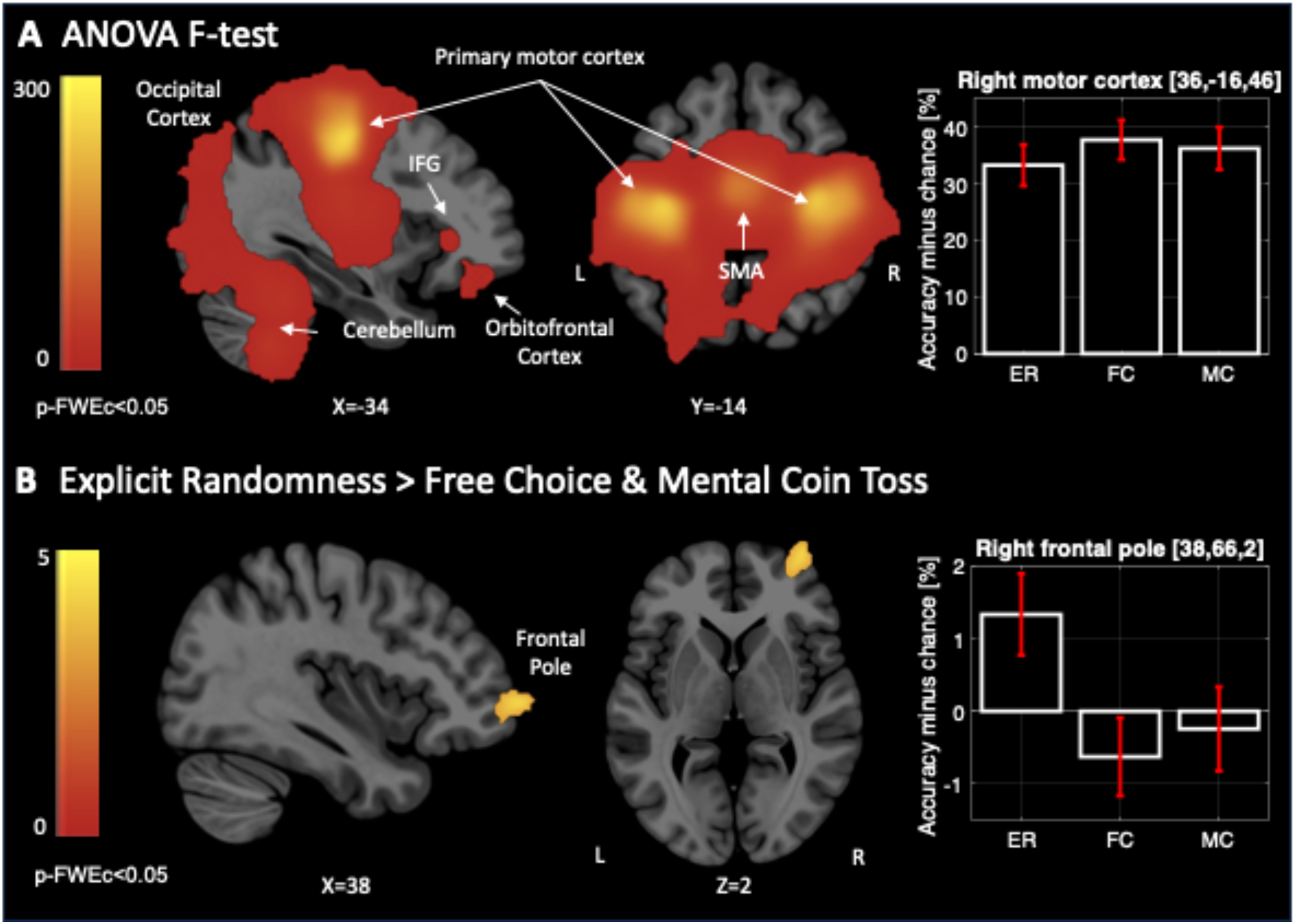
Decoding of trial-by-trial choices using MVPA. Shown are regions where accuracies were above chance level, separately for (A) overall F-test and (b) specific comparison Explicit Randomness > (Mental Coin Toss & Free Choice) overlaid on the Montreal Neurological Institute space (MNI) template (p<0.001, FWE cluster corrected at p<0.05, extent threshold for (A) >=106 voxels and for (B) >=258 voxels). Bar plots show accuracy in percent minus chance level (50%) in right motor cortex (A) and right frontal pole (B), with red error bars indicating the 90% CI. Abbreviations: *IFG*: inferior frontal gyrus, *SMA*: supplementary motor area, *ER*: Explicit Randomness, *FC*: Free Choice, *MC*: Mental Coin Toss.

**Figure 6.**
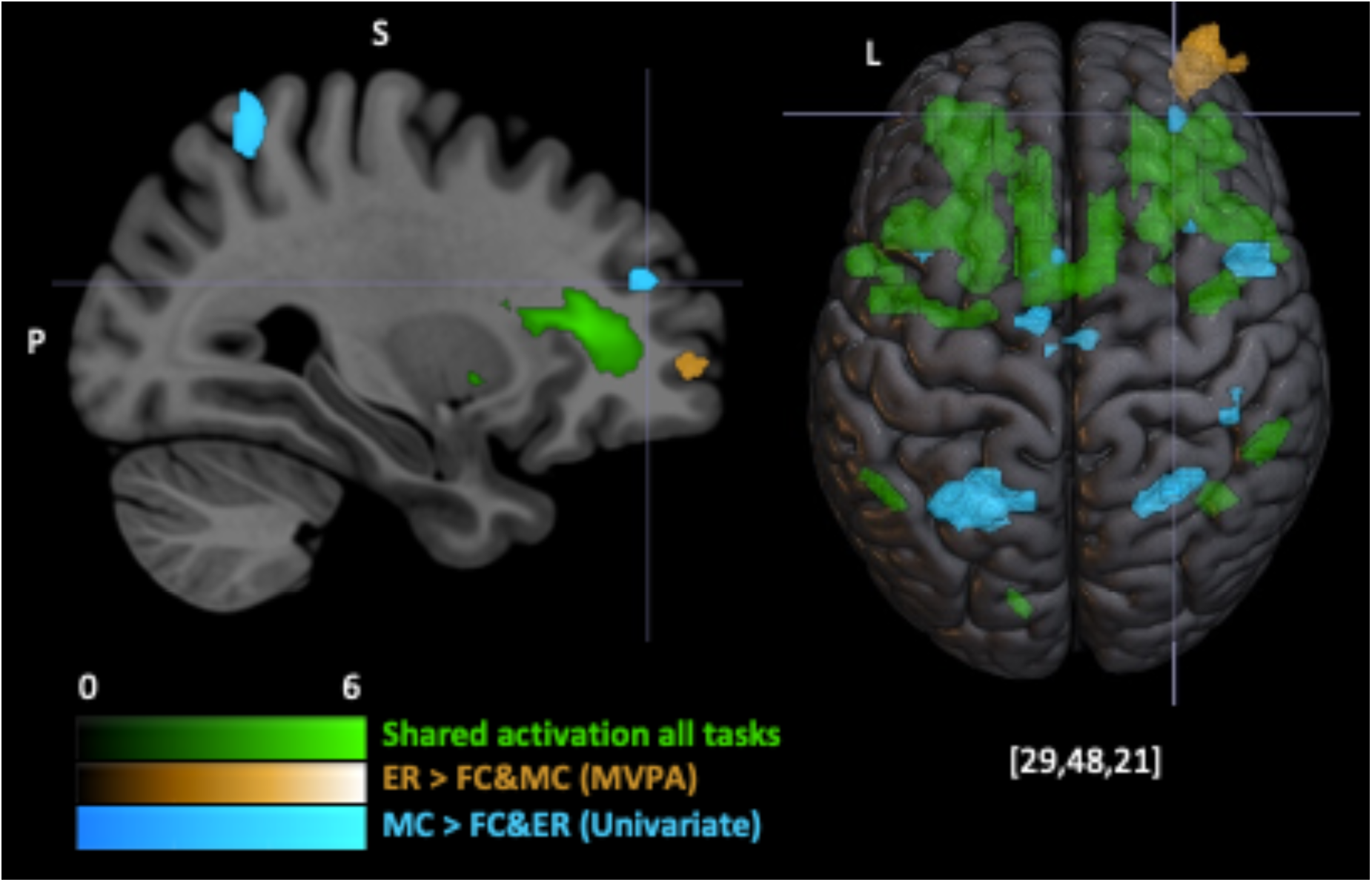
Comparing task-specific and shared activations with informative brain regions in decoding analysis. Results of different contrasts showing differential and shared activations between tasks. Green shows the shared activations across all three tasks as identified via the conjunction analysis. Blue shows the contrast MC>ER&FC in the univariate analysis. Yellow shows the contrast ER>FC&MC from the MVPA analysis. Clusters are overlaid on the Montreal Neurological Institute space (MNI) template, p<0.001, FWE cluster corrected at p<0.05. Abbreviations: *ER*: Explicit Randomness, *FC*: Free Choice, *MC*: Mental Coin Toss.

## Discussion

In this study we explored the neural basis of random sequence generation by giving each subject one of three different instructions for eliciting random behavior. Specifically, participants were asked to conduct a binary decision-making task that instructed them to either (1) make random choices, (2) to make free choices or (3) to choose according to their mental simulation of a fair coin toss. We identified a strong and shared frontoparietal pattern of activity across all three tasks. Our results highlight the significance of mental imagery in the MC task and suggest that regions of the anterior insula (aINS) are distinctly activated in ER and MC as compared to FC. In addition, our MVPA analysis revealed an area in the right frontopolar cortex to be predictive of the choices in the ER task. Our study shows that similar behavioral randomization performance can emerge from distinct cognitive processes.

### Behavioral results

Overall, we observed sequential dependencies in participants’ choice sequences, which is a robust result in the RSG literature (reviewed e.g. by Nickerson, 2002). Drawing from perceptual decision-making research, individuals’ time series exhibit autocorrelations, where current choices are dependent on previous choices. (Urai et al., 2019). Thus, suppression of these autocorrelated responses is not entirely successful, possibly due to constraints posed by the limited capacity controlling mechanism (Jahanshahi et al., 1998). This leads to sequential dependencies in the sequences that were picked up by our metrics.

Notably, all three tasks exhibited a similar randomization profile in terms of our key metrics conditional entropy and Markov order. Our previous study (Guseva et al., 2023), had found that MC sequences differed from those in both ER and FC in terms of conditional entropy, along with differences between MC and FC conditions in terms of Markov orders. Nevertheless, the overall results pattern aligns with Guseva et al. (2023), with MC showing the best randomization performance, followed by ER and then FC in across all metrics. The lack of statistical significance could be attributed to differences in sample size (ranging from 27-29 vs. 73-85 participants per condition in Guseva et al., 2023), sequence length (540 vs. 1000 in Guseva et al., 2023), and trial timing (2000ms vs 1000ms in Guseva et al., 2023).

Despite the absence of explicit randomness cues in the FC instructions, participants’ choices resembled those of the ER and MC randomization tasks, consistent with Naefgen and Janczyk (2018) who found similarities between comparable RSG and FC tasks. The FC task theoretically should not require active memory storage and updating as the choice history should be irrelevant to the task. There is also no apparent need to inhibit habitual or patterned response tendencies, reducing the load on the executive functions. We suggest that participants might have formed certain assumptions about the nature of the task and intuitively introduced randomness into their choices, possibly to give the task a perceived purpose, or perhaps they interpreted a free choice to be a random choice (see post-experiment text responses in our published dataset). This result highlights the role of participants’ assumptions about a cognitive task, as people seem to inject some variability in unconstrained choice situations.

The FC and MC sequences differed in their proportion values, reflecting the balance of heads and tails frequencies. FC participants created less proportional sequences, with one of the sides being chosen more often than the other, consistent with Guseva et al. (2023). This difference may stem from perceiving a coin toss as inherently balanced, e.g., a participant wrote in the post experimental questionnaire: “*At the beginning of the test I thought that after a lot of coin flips the ratio is always around 50-50% between heads and tails, so I tried to press left and right equal times*.“ At the same time, the FC task instructions did not mention any need for balanced choices. In contrast, the average run lengths were comparable across the tasks, showing no significant group differences. This could suggest some inherent pattern length emerging in binary decision scenarios, regardless of instruction.

### Imaging results

#### Common neural substrates across all tasks

We found a large-scale frontoparietal activation pattern across all three tasks. Frontal regions included a bilateral cluster located in IFG, which also extended into regions of aINS and frontal opercular cortex, as well as left pre-SMA, left precentral gyrus/middle frontal gyrus. Parietal regions included bilateral IPL. The cluster also extended into the right cerebellum, left occipital cortex and bilateral putamen. This activation pattern largely overlaps with the frontoparietal activation that is hypothesized to be associated with voluntary decision-making (Rens et al., 2017). Its recruitment could reflect the high demand for cognitive control during all three tasks as will be explained in the following.

The frontal activation pattern comprised a bilateral cluster around the IFG, which is part of the ventrolateral prefrontal cortex (VLPFC). This aligns well with previous findings on RSG (Daniels et al., 2003; Jahanshahi et al., 2000; Koike et al., 2011). However, only little is known about the involvement of IFG in free choice tasks. The robust activation of this area is not unexpected given the established role of VLPFC in action control and executive functions (Segal & Elkana, 2023; Tops & Boksem, 2011). The VLPFC’s executive role is debated, with suggestions of inhibitory control for cognitive and motor interference (Aron et al., 2014; Schaum et al., 2021), and working memory involvement (Segal & Elkana, 2023). Overall, our results highlight the involvement of IFG in randomization tasks.

Of note is the cluster of activation of the posterior portion of IFG in the left hemisphere which corresponds to BA44 (and thus partially overlapping with Broca’s area). This area has not been explicitly observed in either the randomization or free choice literature. While traditionally associated with overt speech production, this region has also been implicated in the generation of internal speech (Alderson-Day & Fernyhough, 2015). This could mean that the participants engaged in internal vocalization of their choices. In fact, some examples of the post-experimental questionnaire contain the following answers: “*Sometimes I focussed [sic] on the choice between left and right by verbalizing it to myself […]*” or “*I did think of the words left/right and the feeling of the movement to either left/right.*” This presence of neural activation in this area opens up the discussion about the role of language-related processes in randomization tasks.

Adjacent to the IFG lies the aINS, which also showed activation across the tasks. In line with our results, Takahashi et al. (2015) found that activity in bilateral insula correlated positively with behavioral entropy during an interactive matching pennies game. Because none of the studies had reported insula activation so far (Daniels et al., 2003; Jahanshahi et al., 2000), the authors concluded that that activation likely arose due to the competitive nature of the task and not due to randomness generation. As our study does not involve a game element, it demonstrates that insula activity can arise in non-competitive RSG task variants nonetheless. Insula activation across all tasks is plausible given its role as a well interconnected integrative hub, involved in cognitive control, goal-directed behavior (Wu et al., 2019), along with motor inhibition (Levy & Wagner, 2011). Together with the IFG, the aINS was suggested to play a crucial role in top-down biasing of the posterior cortex (Tops & Boksem, 2011) reinforcing the role of the frontoparietal network in the three tasks.

We also report activations in left pre-SMA and bilateral clusters in the precentral gyrus, consistent with findings across free choice and the few existing RSG studies. Pre-SMA activity has been associated with higher-level motor planning, internally guided voluntary actions, conflict monitoring and execution of novel actions (Nachev et al., 2007). Our results are in line with the model of corticolimbic control pathways proposed by Tops & Boksem (2011). This framework suggests a chain of top-down control with the IFG/aINS maintaining working memory representations and goals and connection to premotor and motor regions to initiate or suppress responses. This is supported by previous reports of functional connectivity of pre-SMA with the IFG (Schaum et al., 2021; Tomiyama et al., 2022), reinforcing the network’s role in motor response inhibition. We suggest that response inhibition, necessary to override prepotent responses, plays a key role in all three tasks, with premotor along with frontal/insula regions contributing to this process. This offers an alternative neurocognitive possibility compared to Jahanshahi et al.’s (2000) network modulation model.

Our results also revealed shared bilateral activation of the IPL which constitutes part of posterior parietal cortex. This aligns with activations reported in Daniels et al. (2003), while Jahanshahi et al. (2000) found it to be deactivated. IPL activity is also a consistent finding across multiple FC studies (Bode et al., 2013; Lau et al., 2004; Si et al., 2021; Welniarz et al., 2021). Activity in this area might be relevant across all tasks due to its role as a convergence point for multiple networks, contributing to various functions, such as (visuo-) spatial attention and semantic processing (Numssen et al., 2021), attending to current goals and responding to salient information (Singh-Curry & Husain, 2009), mathematical cognition (Wu et al., 2009) and decision-making under uncertainty (Vickery & Jiang, 2009). It is likely that IPL activity is modulated by inferior frontal activation, particularly frontal opercular cortical areas (Higo et al., 2011) positioning it is another pivotal node in top-down control.

Electrophysiological work showed that during a working memory task, prefrontal activity occurs prior to parietal activity, suggesting a hierarchical activation flow with prefrontal areas exerting top-down modulation of parietal regions when implementing cognitive control (Brass et al., 2005). The different task instructions might have activated a host of different goal-directed processes, triggering down-stream effects such as contextual biases on attention processes in more posterior regions (Friedman & Robbins, 2022).

It is unexpected that our analysis did not reveal any joint DLPFC activation across all tasks given the existing literature (Bode et al., 2013; Daniels et al., 2003; Jahanshahi et al., 2000; Lau et al., 2004). DLPFC is a key node in the frontoparietal control network and is strongly connected to the rest of the active regions that were identified in the present analysis, such as IFG, aINS and pre-SMA (Cai et al., 2014; Orr et al., 2015). Jahanshahi et al.’s (2000) neurocognitive model of randomization (network modulation model) specifically highlights the role of the left DLPFC as suppressing prepotent responses by modulating the number association network in superior temporal cortex. Additionally, our initial hypothesis of anterior cingulate cortex activation across all tasks has also not been met, which is surprising as this region appears in tasks with high degrees of decision uncertainty and arbitrary decisions that introduce choice conflicts (Friedman & Robbins, 2022; Furstenberg et al., 2023; Ridderinkhof et al., 2004).

Differences in task setup between the studies might explain the lack of shared activation in both DLPFC and anterior cingulate cortex. Our task differs from Jahanshahi et al. (2000) in that it is a binary choice task and is thus unlikely to induce ascending or descending prepotent responses, as would be the case in a larger alphabetical or numerical choice set, such as digits ranging from 1-9 in Jahanshahi et al. (2000). So, the network modulation model might not be applicable in our case. This raises an important methodological question: Is the randomization across a large, orderly choice set, like the letters of the alphabet merely a variation of the same task or does it represent an entirely different task compared to a binary and/or non-alphanumeric choice set? A clear standardization and taxonomy of randomization tasks is needed. Additionally, in Daniels et al.’s (2003) task the choices were produced faster (1 and 2 Hz) and were generated via an internal speech technique, as opposed to button presses like in our case. Moreover, both studies feature smaller sample sizes (n=8 and n=11).

#### Distinct activations between tasks

Our univariate analysis also compared the activity between the tasks revealing differential activations in aINS, pre-SMA and SPL in both hemispheres. This univariate analysis was based on activation over an entire block, i.e., it captured sustained activation levels during that period. Regions that carried specific choice-predictive information were identified with the decoding approach using single choices. Specifically, we observed the right frontopolar cortex to be predictive of the choices in the ER condition. Interestingly, despite the lack of differences in the behavioral randomization measures, these differential results suggest that participants might have recruited distinct cognitive control mechanisms during the different tasks.

##### Explicit Randomness and Mental Coin Toss

In addition to the shared insular activation, we observed a subregion of the aINS that was more active in both ER and MC as compared to FC. With this region’s role in as a cognitive control hub, the increased activation in ER and MC might reflect a potential higher cognitive load and attention demand in these tasks. The aINS has also been characterized as a bottleneck region of cognitive control capacity (Wu et al., 2019). Alternatively, Jahanshahi et al. (2000) designated the DLPFC as the limited capacity controller. Our results suggest that aINS activation may also play a key role in this capacity, perhaps in tandem with pre-SMA. Overload of the aINS in this subregion might set a limit on randomization performance, potentially explaining why randomization is generally challenging to people. This could occur in the broader context of the cingulo-opercular network where the aINS region is a key node (Menon & D’Esposito, 2022). This difference in activity patterns might indicate that different neural mechanisms are involved in randomization behaviors.

##### Differential activation in Explicit Randomness

The multivariate analysis identified the frontal pole’s most anterior area as predictive of individual choices in the ER condition. Overall this result aligns with what is known about this region as it has been reported to be linked to abstract higher cognition and along multiple functions has been implicated in the evaluation of internally generated information (Christoff & Gabrieli, 2000; Tsujimoto et al., 2010). The frontopolar area has also been associated with monitoring uncertainty and considering alternative courses of action (Wan et al., 2016). The frontopolar activity as observed here could potentially stand atop of the top-down control hierarchy, as it exhibits connections to premotor cortices and parietal lobes (Boorman et al., 2009). At the same time, it is interesting to observe this particular result only in the ER condition, as one might expect these cognitive functions to be shared across all three tasks. It is hence possible that ER might invoke a different cognitive subprocess than a related randomization task, such as MC.

While absent from RSG literature, frontopolar activity has been documented in studies on free choice (Bode et al., 2011; Soon et al., 2008). Interestingly, the observed cluster lies in the vicinity of the region found in Soon et al. (2008), who used a free choice task. While it has been argued that the classifier in Soon et al. (2008) could have captured lingering information from the previous trial due to the autocorrelation present in the choice sequences (Lages & Jaworska, 2012), this was controlled for in subsequent reanalyses (Allefeld et al., 2013). Here this possibility was ruled out, as such a result would have been observed in all conditions, given that all conditions exhibit some degree of autocorrelation not just the ER.

It is worth noting that in Soon et al. (2008), subjects were pre-selected who already produced balanced sequences, which could have effectively reduced the dataset to choice sequences resembling those from an ER task rather than a true FC task. As we outlined above, a free choice might be understood as a random choice by people or at least carry this connotation because it is common practice to include specific restrictions and rules (e.g., to avoid repetitions) in “free choice” instructions (Naefgen & Janczyk, 2018). In contrast, our FC instructions were intentionally unrestricted. This specific type of FC task instructions lead to a differential neural activation profile compared to a simple randomization task, challenging the contemporary view that ER and FC are the same task (Naefgen & Janczyk, 2018). For future experiments it would be advisable to have better ways to define what an FC task is and to closely examine how people understand it.

##### Differential activation in Mental Coin Toss

The MC task activated frontoparietal regions stronger than the other two tasks. The key regions in the frontal lobe that showed heightened differential activation were pre-SMA/SMA, IFG and DLPFC. We also observed bilateral activation of SPL. This activation pattern strongly resembles the network engaged during visuo-spatial motor imagery, encompassing the IFG, SMA, aINS as well as inferior and superior parietal lobules. Also, subcortical regions such as cerebellum and thalamus have been implicated in motor imagery (Hétu et al., 2013). This is consistent with our expectation that a mental coin toss task likely involves simulating motor actions. Martel and Glover (2023) argue that motor imagery requires executive resources that are mediated by DLPFC activation. This could potentially explain the specific dorsolateral activation in MC and not in the other condition.

The differential involvement of the right IPS also aligns with the task demands as it has been involved in visuo-spatial attention and working memory (Bray et al., 2015). Notably, with the similar behavioral performance across tasks, this suggests that perhaps motor imagery alone does not seem to pose a significant randomization advantage compared to the other tasks.

On the surface, it seems that our instruction conditions produce a similar outcome. However, “underneath the hood” they partially activate different neural pathways. Considering the pervasive issue of differing task parameters (such as task instructions) across randomization experiments coupled with the lack of standardized experimental protocols, and taking into account our observation that instructions matter on a neural level, what has truly been measured in the past decades of research on RSG? Our results suggest that it is important to take a step back and build a stronger methodological foundation by re-examining the most basic components of the RSG task.

### Limitations

Several limitations of this study should be considered. First, the primary methodological difficulty is in assessing performance in tasks where choices are generated internally (as opposed to stimulus-driven), lacking a clear objective criterion to confirm whether participants actually correctly understood and executed the task. Unlike in perceptual decision-making research for instance, where accuracy scores allow the researcher to evaluate (1) compliance and/or (2) correct understanding of the task, this is challenging in tasks without a criterion for right or wrong answers. For instance, an important factor specifically in randomization tasks, is participants’ familiarity with random processes. Differences in statistical literacy and their impact on randomization are still unexplored.

Thus, to gauge participants’ understanding and attention to the task we used questions and open-form text responses. As this is a rather simple approach, we suggest the field may benefit from developing a robust method to closely track and unambiguously assess participants’ understanding and adherence to instructions.

Second, we used established randomness measures that are sound and grounded in theory. Nonetheless, there is no measure that can comprehensively test every aspect of randomness in a sequence (Ayton et al., 1989), especially considering that human-made sequences are naturally rather short. Therefore, the absence of performance differences in this study does not rule out the possibility that other measures might identify them.

Finally, although we identified basic neural substrates of randomization, the specific roles of the neural regions remain speculative. Further investigation, for instance within the context of Cooper’s (2016) computational model of randomization, could provide a deeper understanding of the circuits supporting various cognitive functions.

### Conclusion

Our exploratory study into the neurocognitive architecture of randomization across different task instructions found consistent frontoparietal pattern of activation across all three tasks, indicating shared neural substrates associated with cognitive control and inhibitory processes. Some of our results differ from the few previous studies in terms of dorsolateral PFC and anterior cingulate cortex activation, possibly due to increased power with a substantially enlarged sample. In addition to the shared activation, our findings also suggest that although the three tasks produce similar behavioral outcomes, they also partially rely on distinct neural pathways. Our results highlight the complexity of brain-behavior relationships (Gilman et al., 2015) and challenge the idea of a universal random generator in the brain.

## Data availability statement

The functional images are available upon request. The code for both behavioral and fMRI analysis as well as the behavioral dataset are available here: https://github.com/m-guseva/RANDECOD_fMRI_study.

## Author contribution

- Maja Guseva: Conceptualization, Data Curation, Formal Analysis, Investigation, Methodology, Project Administration, Resources, Software, Validation, Visualization, Writing – Original Draft Preparation, Writing – Review & Editing
- Carsten Bogler: Conceptualization, Methodology, Supervision, Validation, Writing – Review & Editing
- Carsten Allefeld: Methodology, Writing – Review & Editing
- Ece Büşra Ziya: Investigation
- John-Dylan Haynes: Conceptualization, Methodology, Funding Acquisition, Resources, Supervision, Validation, Writing – Review & Editing

## Acknowledgements

We have no conflicts of interest to disclose. Correspondence concerning this article should be addressed to Maja Guseva, Bernstein Center for Computational Neuroscience, Charité Universitätsmedizin Berlin, Charitéplatz 1, 10117 Berlin, Germany.

## Citation Diversity statement

Proportion of categorized DOIs by category: M/M: 0.603, W/M: 0.224, M/W: 0.155, W/W: 0.017

Gender Citation Balance Indices (GCBIs) by category: M/M: 0.483, W/M: -0.3, M/W: 0.349, W/W: -0.892

